# A Multidimensional Analysis of the Bimodal Piperaquine Response in *Plasmodium falciparum*

**DOI:** 10.64898/2026.07.06.734584

**Authors:** John Kane, Aubrey Schall, Lisa A. Checkley, Douglas A. Shoue, Sydney M. Gavula, Caroline Thomas, Xue Li, Ian H. Cheeseman, Ashley M. Vaughan, Timothy J. Anderson, Manuel Llinás, Paul D. Roepe, Michael T. Ferdig

## Abstract

Malaria remains a pressing global health challenge, with the continued emergence of resistance threatening the long-term efficacy of artemisinin-based combination therapies (ACTs). Piperaquine (PPQ), an important partner drug in artemisinin-based combination therapies exhibits a unique bimodal dose-response phenotype associated with reduced susceptibility, yet the biological mechanism underlying this phenotype remains unknown. This phenotype is strongly associated with mutations in pfcrt and copy number amplification of *plasmepsin II/III* (*pm II/III*). Given that plasmepsins play a central role in hemoglobin degradation within the blood stage parasite digestive vacuole, and that PPQ accumulates within this compartment and perturbs heme detoxification, this phenotype likely reflects alterations in fundamental biological processes alongside drug-specific effects. We used isogenic PPQ-resistant parasite clones differing only in *pm II/III* copy number to integrate phenotypes with metabolic changes, and transcriptional responses to ascertain the impact of genotype combinations on parasite response to PPQ. Across increasing PPQ concentrations, parasites with elevated *pm II/III* copy number exhibited distinct metabolic responses compared to single-copy parasites, specifically, an altered abundance of peptides derived from hemoglobin degradation, directly implicating a core biological pathway long associated with plasmepsin function. The combination of metabolic and transcriptional data with phenotypic measurements supports a model in which increased plasmepsin expression enhances the parasite’s capacity to sustain hemoglobin digestion and associated metabolic activity under high PPQ concentrations. This points to a mechanistic basis for continued parasite survival, indicating that changes in hemoglobin processing within the digestive vacuole contribute to the bimodal response to PPQ. Molecular dynamics simulations further support a direct interaction between PPQ and PM II/III, as a mechanism by which these proteins impact PPQ response dynamics through both modulation of hemoglobin digestion and protein-drug interactions within the digestive vacuole.

## Introduction

With over 250 million cases reported in 2024, malaria remains a major global health burden (1). Despite substantial reductions in morbidity and mortality over the past two decades, the continued emergence of drug-resistant *Plasmodium falciparum* represents the greatest threat to control and eradication efforts. Artemisinin-based combination therapies (ACTs) are used globally as the current frontline treatment for uncomplicated malaria. However, reduced susceptibility to artemisinin, associated with mutations in *kelch13*, first established in Southeast Asia, is increasingly being observed in Africa, where delayed parasite clearance and the emergence of novel *kelch13* variants raise intense concerns about the future efficacy of these therapies (2,3). The reduced efficacy of artemisinin, the rapidly acting component of ACTs, places a larger burden on the slower-acting partner drugs to clear the remaining parasite biomass. This shift in therapeutic dependence highlights the need to define the molecular mechanisms that confer resistance to frontline ACT partner drugs. Piperaquine (PPQ) remains an important ACT partner drug, as part of the DHA+PPQ combination therapy, and is deployed in several malaria-endemic regions as a second-line therapeutic (1). However, the emergence of PPQ treatment failure in Southeast Asia threatens the long-term efficacy of this important ACT partner drug (2,4–7). Despite the identification of multiple genetic associations with reduced PPQ susceptibility, the mechanistic basis by which these variants contribute to the resistance phenotype remains poorly understood.

Genetic studies of PPQ-resistant parasites have identified mutations in the *chloroquine resistance transporter* (*pfcrt*) and copy number amplification of *plasmepsin II* and *plasmepsin III* (*pm II/III*), which encode digestive vacuole proteases involved in the initial steps of hemoglobin (Hb) degradation, as major genetic variants associated with reduced PPQ susceptibility (4,6,8,9). Two independent genetic crosses between Cambodian PPQ-resistant parasites and African PPQ-sensitive lines further resolved the relative contributions of these resistance loci, demonstrating that *pfcrt* is the primary determinant of PPQ resistance, while the influence of *pm II/III* copy number amplification is largely restricted to parasites carrying a permissive PPQ-resistant pfcrt background (10,11). Additional loci, including mutations in an exonuclease (*pfexo*) and *pfatg18*, also have been associated with the PPQ response, though their functional roles are poorly understood (9–11). Recombinant progeny from these crosses further show that variation in *pm II/III* copy number has only a modest effect in the absence of a permissive *pfcrt* background and is dependent on the specific phenotype measured (10). Consistent with these findings, targeted genetic studies demonstrated that introduction of *pfcrt* resistance mutations into chloroquine-resistant laboratory strains, including Dd2 and GB4, is sufficient to confer reduced PPQ susceptibility (6,12,13), whereas manipulation of *pm II/III* copy number has more limited and context-dependent effects, with gene disruption increasing dose susceptibility but overexpression alone failing to recapitulate the full resistance phenotype (14,15).

In contrast to most antimalarial drugs, where *in vitro* resistance is characterized by a progressive rightward shift in the dose-response curve and an increase in IC_50_, PPQ resistance does not follow this conventional profile. Instead of a sigmoidal dose-response curve, most PPQ-resistant parasites exhibit a bimodal response, characterized by a secondary phase of increased survival at high drug concentrations (16). This non-canonical phenotype has been strongly associated with copy number amplification of *pm II/III*, as parasites carrying a single copy display a typical sigmoidal response, whereas those with multiple copies develop a pronounced secondary survival peak, the magnitude of which increases with copy number (16). Supporting a functional role for these proteases, knockout of *pm II/III* reduces the magnitude of the secondary survival phase, with single *pm III* and double *pm II/III* knockouts abolishing bimodality entirely, whereas *pm II* knockout results in a substantial but incomplete reduction of the high-dose survival peak (17). Consistent with the broader epistatic effects observed for *in vitro* PPQ resistance using classical genetic approaches, this phenotype is not driven by *pm II/III* amplification alone. Within our previous genetic cross, recombinant progeny inheriting the drug-sensitive 3D7-like *pfcrt* allele do not exhibit a bimodal dose-response regardless of *pm II/III* copy number, demonstrating that this phenotype is restricted to parasites carrying PPQ-resistant *pfcrt* alleles (10,11).

Beyond its novel presentation, the bimodal PPQ dose-response complicates standard approaches for measuring *in vitro* drug susceptibility. The presence of a secondary survival phase prevents accurate estimation of IC_50_ values using conventional curve-fitting methods, as high-dose survival is not captured by a single sigmoidal model. As a result, alternative metrics have been adopted, including quantification of the area under the curve (AUC) to capture the magnitude of the secondary response (16), as well as the piperaquine survival assay (PSA), which measures parasite survival at clinically relevant drug concentrations that fall within this high-dose survival range (18).

While the genetic factors associated with PPQ resistance and bimodal dose-response phenotype have been identified, the mechanistic basis underlying this phenotype is unresolved. Here, we take a comprehensive approach to define the biological basis of this phenotype. We show that the bimodal PPQ survival pattern is stage-specific across the erythrocytic developmental cycle and is shaped by the timing of drug exposure, indicating that this response is driven by interactions and processes upon initial PPQ exposure. In parallel, increased *pm II/III* copy number is associated with substantial shifts in the abundance of Hb-derived peptides under increasing PPQ concentrations, reflecting differences in the parasite’s metabolic state under drug pressure. Molecular dynamics simulations further identified a favorable interaction between PPQ and PM II/III, supporting a model in which increased plasmepsin abundance increases the capacity for PPQ-plasmepsin interactions during drug exposure. Together, these findings support a new mechanistic model in which excess PM II/III abundance from genomic copy number amplification enables these proteins to interact with PPQ while maintaining sufficient proteolytic activity to sustain hemoglobin metabolism, thereby promoting parasite survival under high-dose PPQ exposure.

## Results

### Stage specificity of the PPQ dose-response curve shape

To determine whether parasite stage influences the bimodal survival response to PPQ, tightly synchronized parasites were subjected to a 12-point dose-response assay following initial drug exposure at defined stages of the erythrocytic developmental cycle. We compared a PPQ-resistant line with 4 copies of *pm II/III* exhibiting a bimodal response (KH004), an isogenic single-copy *pm II/III* derivative with a sigmoidal response (FG0305), and a drug-sensitive parasite with a single copy of *pm II/III* (MAL31) with a sigmoidal response. Resistant parasites KH004 and FG0305 both carry the novel Dd2+G367C *pfcrt* genotype, whereas MAL31 carries a PPQ-sensitive 3D7-like genotype (10).

While both *pm II/III* single-copy parasites generated consistent sigmoidal dose-response curves regardless of stage, KH004 exhibited a pronounced stage-dependent phenotype (Fig. 1A-C). Specifically, KH004 lost the bimodal nature of its dose-response curve when first exposed to PPQ as schizonts, whereas exposure at ring and trophozoite stages preserved the bimodal response (Fig. 1A). This pattern was also observed in genetically distinct Southeast Asian isolates carrying PPQ-resistant *pfcrt* alleles and multiple copies of *pm II/III* as well as recombinant progeny from a genetic cross between KH004 and MAL31.

**Fig. 1.**
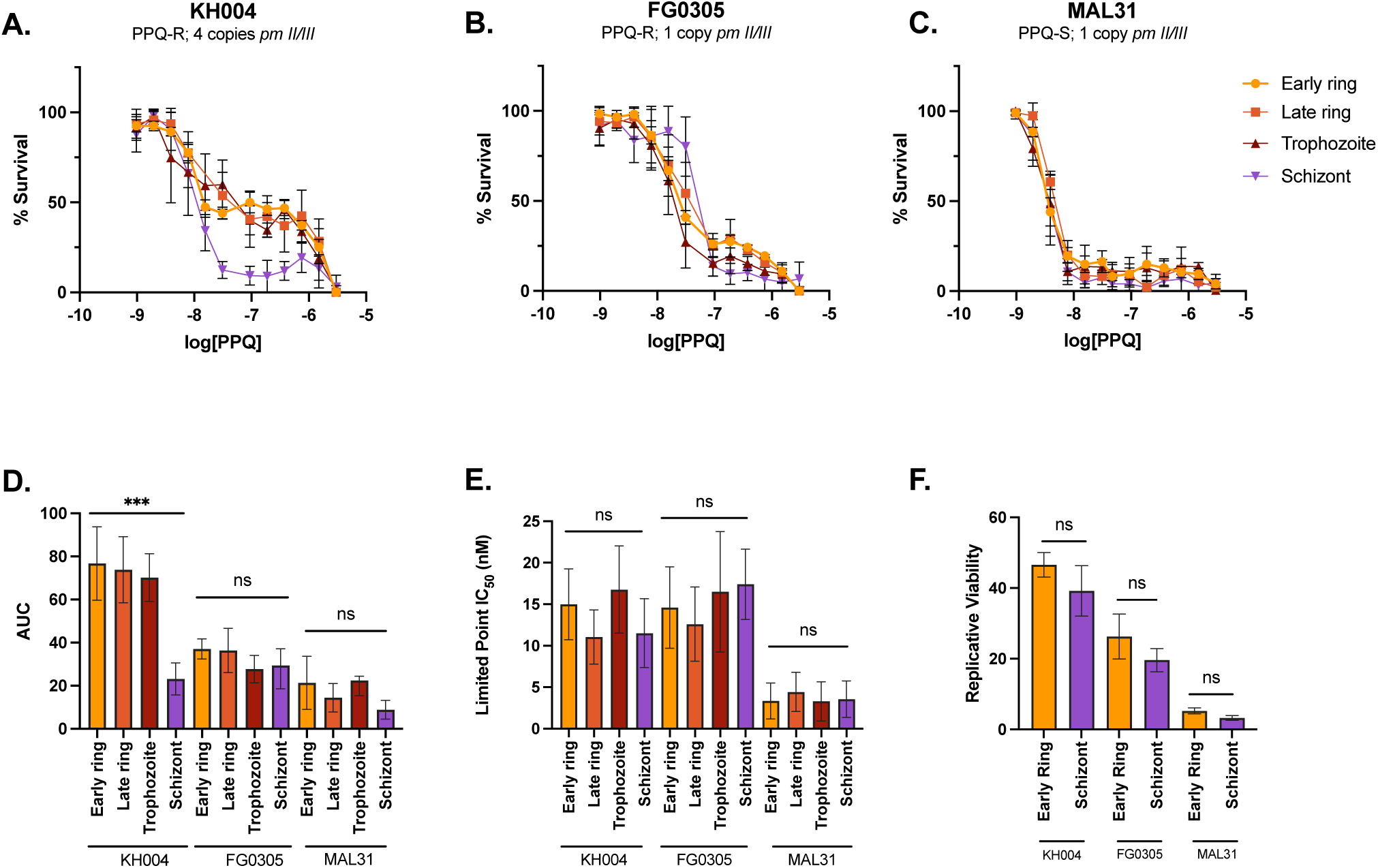
Stage specificity of bimodal dose-response curve. A-C) Parasites with varying combinations of *pm II/III* copy number and *pfcrt* allele were subjected to standard 72h drug susceptibility assays with varying stages of initial exposure. Stage had no impact on single copy *pm II/III* parasites, regardless of overall PPQ resistance level, but impacted parasites with multiple copies of *pm II/III*. KH004 responded similarly between ring and trophozoite stages, producing a bimodal curve, but lost the secondary phase of the curve when initially exposed at schizont stage. D) This change is quantified through significant reduction in AUC in KH004. E-F) Despite the impact on shape of the dose response curve, PPQ susceptibility in general was not found to be stage specific, either through limited point IC50 or PSA. Statistical significance was determined via one-way ANOVA and Mann-Whitney U tests, and bar graphs represent mean +/- SEM.

The change in curve shape was quantified through AUC of the region encompassing the secondary phase in the bimodal curve (Fig. 1D). To assess the role of parasite stage on overall susceptibility to PPQ, we also calculated limited-point IC_50_ and performed PSA at the all parasite stages (Fig. 1E-F). No significant change in resistance was noted using either IC_50_ or PSA when comparing between initial exposure stages. These results indicate that the bimodal survival phenotype is stage-dependent and that the contribution of *pm II/III* copy number to this phenotype is distinct from the broader PPQ resistance phenotype associated with *pfcrt*.

### Duration of PPQ exposure does not influence the presence of bimodal survival phenotype

The standard 72-hour dose-response assay used to quantify PPQ susceptibility spans approximately 1.5 *P. falciparum* asexual blood stage cycles of, resulting in unequal representation of intraerythrocytic stages during drug exposure. Given the stage specificity of the bimodal survival phenotype (Fig. 1), we sought to determine whether this phenotypic response was driven by the stage at initial exposure or by cumulative effects of prolonged drug pressure over the duration of the assay.

To address this, PPQ exposure was limited to 24- or 12-hour windows, after which parasites were returned to drug-free media for the remainder of the 72-hour assay. These reduced-exposure experiments were initiated with both tightly synchronized ring- and schizont-stage parasites. As expected for a slow-acting antimalarial like PPQ, shorter exposure resulted in a rightward shift in dose-response curves and increase in IC_50_ values. However, the overall shape of the dose-response curve remained unchanged (Fig. 2). Ring-stage KH004 parasites retained a bimodal response even when PPQ exposure was restricted to a 12-hour window entirely within the ring stage, indicating that this phenotype does not require transitions between developmental stages (Fig. 2A). In contrast, parasites initially exposed as schizonts maintained a sigmoidal dose-response regardless of exposure duration (Fig. 2B). Together, these findings indicate that the bimodal survival phenotype is determined by the stage at initial PPQ exposure rather than by cumulative drug exposure across developmental stages.

**Fig. 2.**
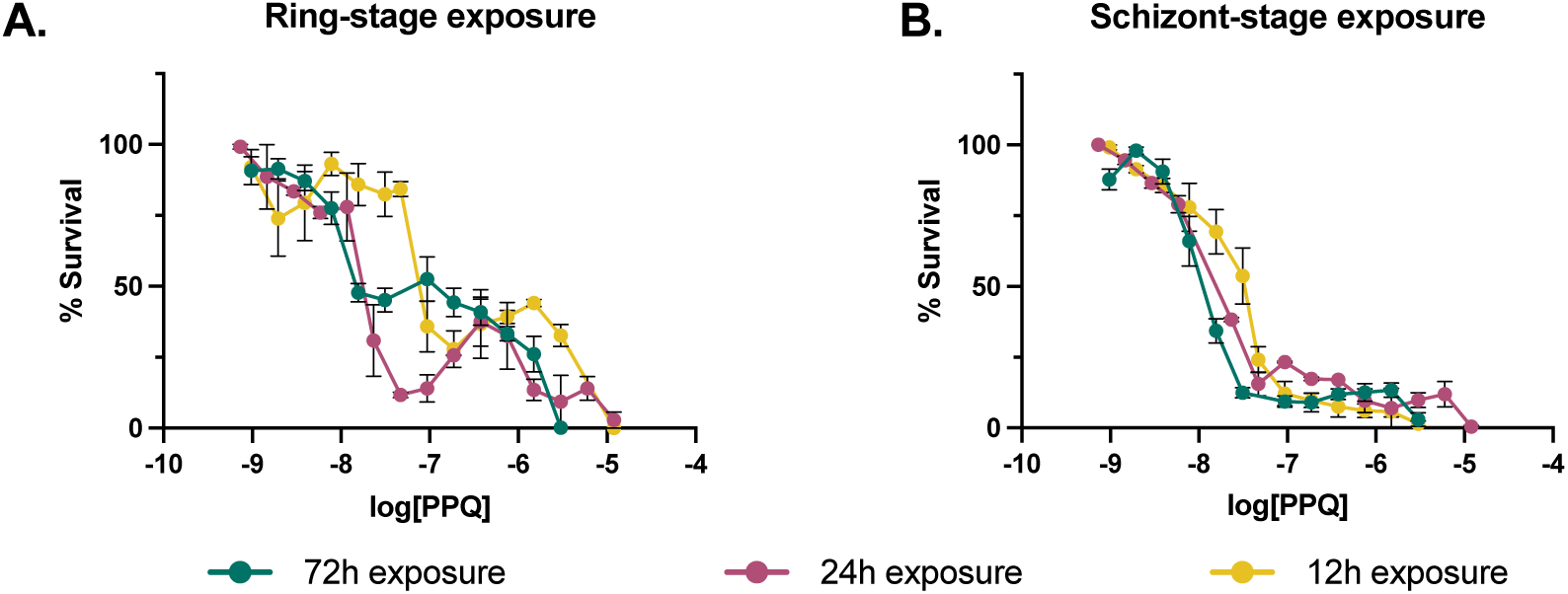
Impact of decreased exposure time on KH004 dose-response shape. Compared to a standardized drug susceptibility assay of 72h of continuous drug exposure, exposure time was reduced to either 24h or 12h, with an outgrowth in untreated complete media for the remainder of the 72h. Due to the slower mechanism of action of PPQ, reduced exposure time shifted the IC_50_ upwards. However, reduced exposure time did not impact the overall bimodal nature of the KH004 dose response curve in ring stage parasites. Similarly, reduced exposure time did not recover the bimodal curve shape in KH004 parasites exposed as schizonts, indicating that this loss of bimodality is the result of interaction between schizont stage parasites and PPQ. This trend of shifting the IC_50_ without changing curve dynamics held true for all parasites, regardless of resistance status or *pm II/III* copy number.

### Exposure to PPQ does not induce overexpression of *pm II/III*

To determine whether inducible changes in *pm II/III* expression contribute to the bimodal PPQ survival phenotype, we quantified transcript abundance across the parasite lifecycle and following PPQ exposure. Because PPQ-resistant parasites remain susceptible at moderate drug concentrations but exhibit substantial survival at higher concentrations, we hypothesized that elevated PPQ exposure may trigger increased *pm II/III* expression under conditions associated with the secondary survival phase.

Consistent with prior observations (19), *pm II/III* expression followed a cyclical pattern across the asexual lifecycle, with peak expression during the trophozoite stage and markedly reduced expression during schizogony. We observed the same stage-dependent expression profile across parasites regardless of underlying *pm II/III* copy number, although overall transcript abundance scaled with copy number (Fig. S1). These data indicate that increased *pm II/III* copy number elevates overall transcript abundance, which has been previously noted in genetically modified parasites (20), while remaining constrained by normal developmental regulation. Notably, these findings suggest that the phenotypic consequences of *pm II/III* amplification depend not only on genomic copy number but also on stage-specific transcription. During schizogony, transcription declines to such low levels that the expression advantage conferred by gene amplification is largely lost, potentially explaining the disappearance of the bimodal survival phenotype.

Although increased *pm II/III* copy number explains differences in PPQ susceptibility between parasite lines, it does not explain the paradoxical increase in parasite survival observed at high PPQ concentrations relative to intermediate concentrations. We next tested whether increasing PPQ exposure alters *pm II/III* expression in isogenic parasites KH004 and FG0305 following short-term exposure to either 50 nM or 200 nM PPQ. These concentrations were selected to capture conditions under which both parasites exhibit similar susceptibility (50 nM) or where KH004 displays increased survival while FG0305 remains inhibited (200 nM) (Fig. 3A). Parasites were exposed for 2h prior to transcript quantification, a time window sufficient to detect transcriptional responses while minimizing stage-dependent variation.

**Fig. 3.**
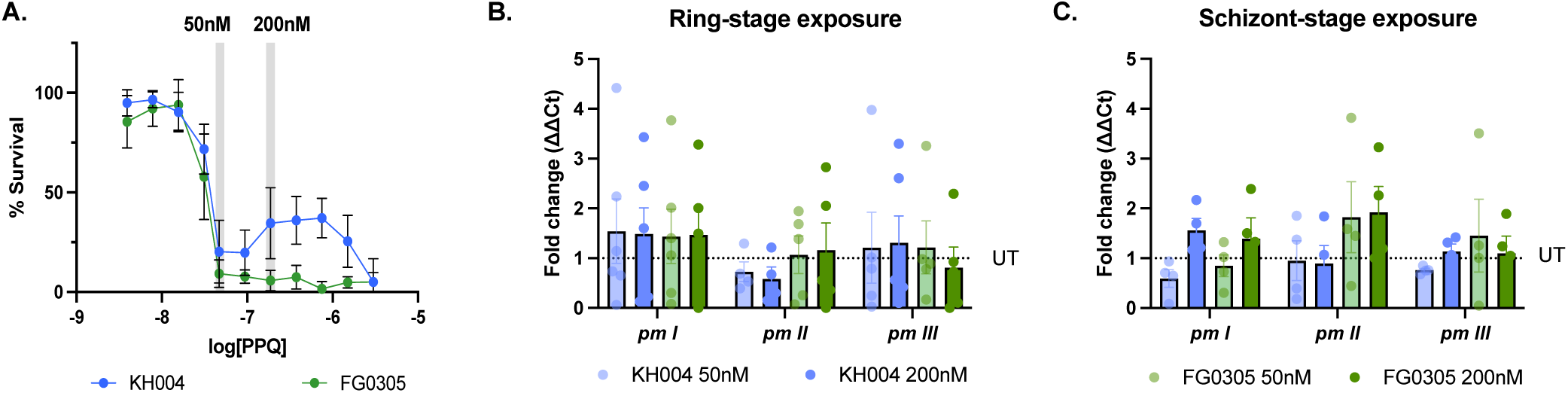
Transcriptional response of *pm II/III* to PPQ exposure. Isogenic parasites KH004 (4 copies *pm II/III*) and FG0305 (1 copy *pm II/III*) were used for assessing changes in *pm I-III* expression under PPQ pressure. **A)** Transcript abundance was measured following a 2h exposure of either no drug, 50nM PPQ, or 200nM PPQ, which corresponds to unique parts of the dose response curve. **B)** This analysis was performed on ring stage parasites, where KH004 has a bimodal curve and FG0305 does not, as well as **C)** schizont stage, where both parasites have identical sigmoid dose-response phenotypes. Compared to untreated samples (dotted line), we do not observe any significant changes in *pm I-III* transcript abundance in the presence of PPQ in either ring stage or schizont stage parasites. The region of *pm I* used for assessing transcript abundance is not part of the gene duplication, therefore, both KH004 and FG0305 have a single copy of this gene. All samples were normalized to *actin1* and compared to untreated condition for the same parasite. Bars represent average fold change of ΔΔCt values (Treated:UT) +/- SEM.

Under these conditions, no significant changes in *pm II/III* expression were observed in response to increasing PPQ concentrations relative to untreated controls in either parasite line (Fig. 3). The transcriptional response to PPQ exposure was similarly unchanged in both ring- and schizont-stage parasites, indicating that drug-induced variation in *pm II/III* expression does not differ across developmental stages. Together, these results indicate that the increased survival observed at high PPQ concentrations is not driven by inducible overexpression of *pm II/III*.

### *pm II/III* copy number alters abundance of Hb-derived peptides under high-dose PPQ

Because PPQ-induced changes in *pm II/III* expression did not explain the concentration-dependent bimodal survival phenotype, we next investigated whether increased *pm II/III* copy number alters parasite metabolism, given the central role of PM II/III in Hb degradation (21–23). Using targeted metabolomic analyses on isogenic PPQ-resistant parasites we evaluated the effect of increasing *pm II/III* copy number on parasite metabolic responses under PPQ conditions associated with the bimodal phenotype.

Three isogenic KH004-derived parasite lines carrying one, three, or five copies of *pm II/III* were profiled under five treatment conditions selected to capture distinct phenotypic outcomes across a range of PPQ exposures. The 50 nM and 200 nM treatment conditions were chosen to align the metabolomic analyses with the transcriptional experiments. Previous research has demonstrated that treatment at high drug concentrations, typically 10× IC_50_, produces the most robust and reproducible metabolomic signatures across diverse antimalarial compounds (24). Therefore, to maintain continuity with the transcriptional experiments while optimizing metabolite detection, we included proportionally scaled PPQ concentrations of 500 nM and 2 µM, corresponding to 10× the 50 nM and 200 nM transcriptional treatment conditions. An additional 200 nM PPQ condition was included to preserve a direct comparison with the transcriptional analyses. Untreated parasites and atovaquone (ATQ; 10 nM, 10× IC_50_) served as negative and positive controls, respectively.

Following metabolite extraction and analysis by liquid chromatography mass spectrometry (LC-MS), 123 targeted metabolites passed quality control filtering and were included in downstream analyses. As expected, ATQ exposure resulted in increased abundance of N-carbamoyl-L-aspartate a characteristic metabolic consequence of cytochrome bc1 inhibition and the resulting impairment of DHODH-dependent pyrimidine biosynthesis (24), confirming metabolic activity across all three parasite lines (Fig. S2).

Exposure to 2 µM PPQ produced *pm II/III* genotype-dependent metabolic differences across the isogenic parasite lines. In the single-copy line FG0305, only one of the 123 targeted metabolites, the Hb-derived peptide PVNF, increased significantly relative to untreated parasites (Fig. 4A). In contrast, parasites harboring multiple copies of *pm II/III* exhibited broader increases in Hb-derived peptides following 2 µM PPQ exposure, with the number of significantly altered peptides increasing with copy number. The three-copy KH004-E9 clone showed increased abundance of PE, DLH, and VD, while the five-copy KH004-D9 clone displayed increases in PE, PD, DLH, VD, and PVNF (Fig. 4 B-C). These data indicate that higher *pm II/III* copy number is associated with greater accumulation of Hb-derived peptides under high-dose PPQ exposure, consistent with altered Hb metabolism in parasites that exhibit increased survival at this concentration. By contrast, exposure to 200 nM or 500 nM PPQ resulted in few metabolites reaching significance relative to untreated controls (Fig. S3), consistent with previous peptidomics-based studies using similar low concentrations of PPQ (17).

**Fig. 4.**
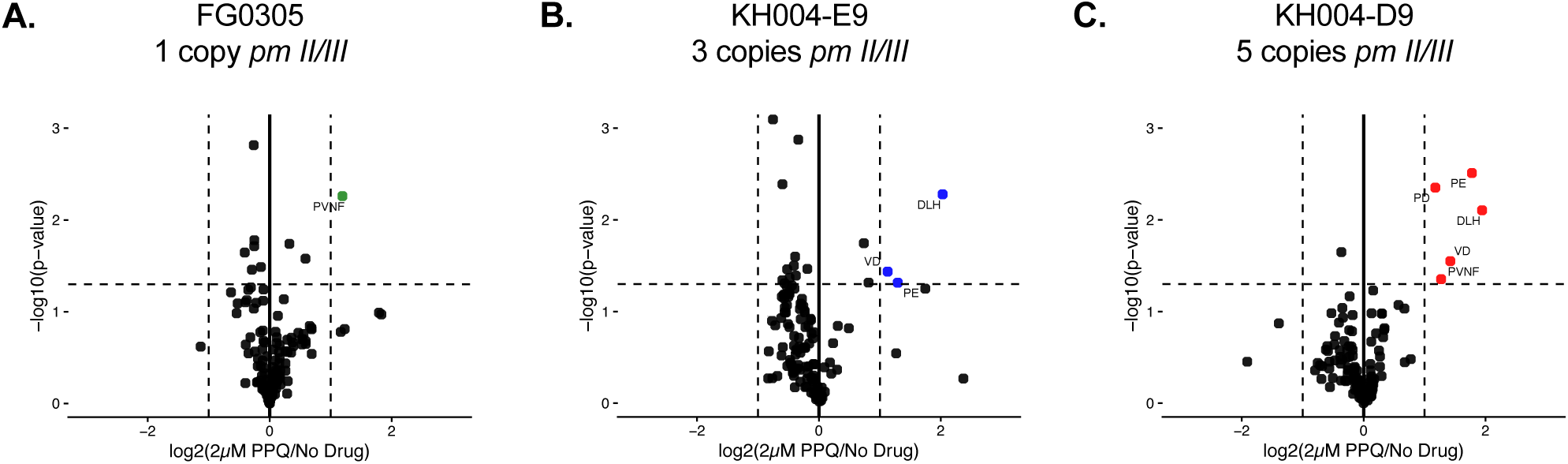
Increased *pm II/III* copy number leads to increased abundance of hemoglobin derived metabolite under high-dose PPQ exposure. Volcano plots showing differences in metabolite abundance relative to untreated controls following exposure to 2 µM PPQ. Columns from left to right correspond to FG0305 (1 copy of *pm II/III*), KH004-E9 clone (3 copies of *pm II/III*), and KH004-D9 clone (5 copies of *pm II/III*). Each point represents a detected metabolite plotted as log_2_ fold change relative to untreated parasites (x-axis) versus –log_10_ (p-value) (y-axis), with colored and labeled metabolites reaching statistical significance. Vertical dashed lines indicate fold-change thresholds and the horizontal dashed line indicates the significance threshold.

### Molecular docking of PPQ to PM II/III

Finally, because PM II/III are digestive vacuole proteases rather than canonical drug transporters or established antimalarial targets, we investigated whether their contribution to the bimodal PPQ phenotype could involve direct interaction with PPQ. To evaluate this, molecular docking simulations were performed using energy-minimized structures of the mature enzymes (sequences corresponding to the enzyme with N terminal prodomain removed) and PPQ ligand. PM II XR (x-ray crystal structure; 1LF3 | pdb_00001lf3) and AF2 (AlphaFold 2 AI structure) and PM III AF2 pdb structures underwent replica exchange molecular dynamics (REMD) minimization three times, each simulation running for 100ns (3×100ns), with 10 ps time steps, at pH=5.0, temperature=310°K, and the protein solvated in simple point charge water and ions added to counter charge as needed, as described earlier for PfCRT protein (25). The second half of frames from the simulation that corresponded to time after the solved structure had reached plateau were then averaged together to create the molecular dynamics (MD) minimized structures for the proteins (AFMD, XRMD). All atom RMSD of the energy minimized PMII x-ray and AF2 structures was 2.36 Å (Fig. S4), providing additional confidence for the veracity of the AF2 structures.

For PM II, subsequent docking of this minimized structure to PPQ identified two binding sites (Fig. 5). The highest affinity pose for each site was energy minimized in the same way as for the undocked proteins described except it was run with 100 ps time steps and all frames were used to acquire the averaged structure. The primary site (site 1) corresponded to the catalytic cleft, where PPQ was positioned between catalytic dyad residues Asp34 and Asp214 (Fig. 5 A-B). A secondary site (5/27 total poses from three independent docking simulations) was observed within a pocket formed by the Loop 3 (L3) and Loop 4 (L4) regions of the protein, adjacent to Leu242 and Gln275, residues previously implicated in substrate binding during Hb cleavage (26) (Fig. 5 D-E). When examining the three highest affinity poses for each of three independent docking trials, PPQ localized to the catalytic site in 8 of the 9 highest affinities poses, with 1 occupying the L3–L4 site 2 pocket (Table 1). By comparison, Pepstatin A, a well-characterized aspartic protease inhibitor, localized exclusively to the enzyme’s active site, consistent with its established mode of inhibition (Fig. S5). Notably, the catalytic region corresponds to site 1 occupied by PPQ.

**Figure 5.**
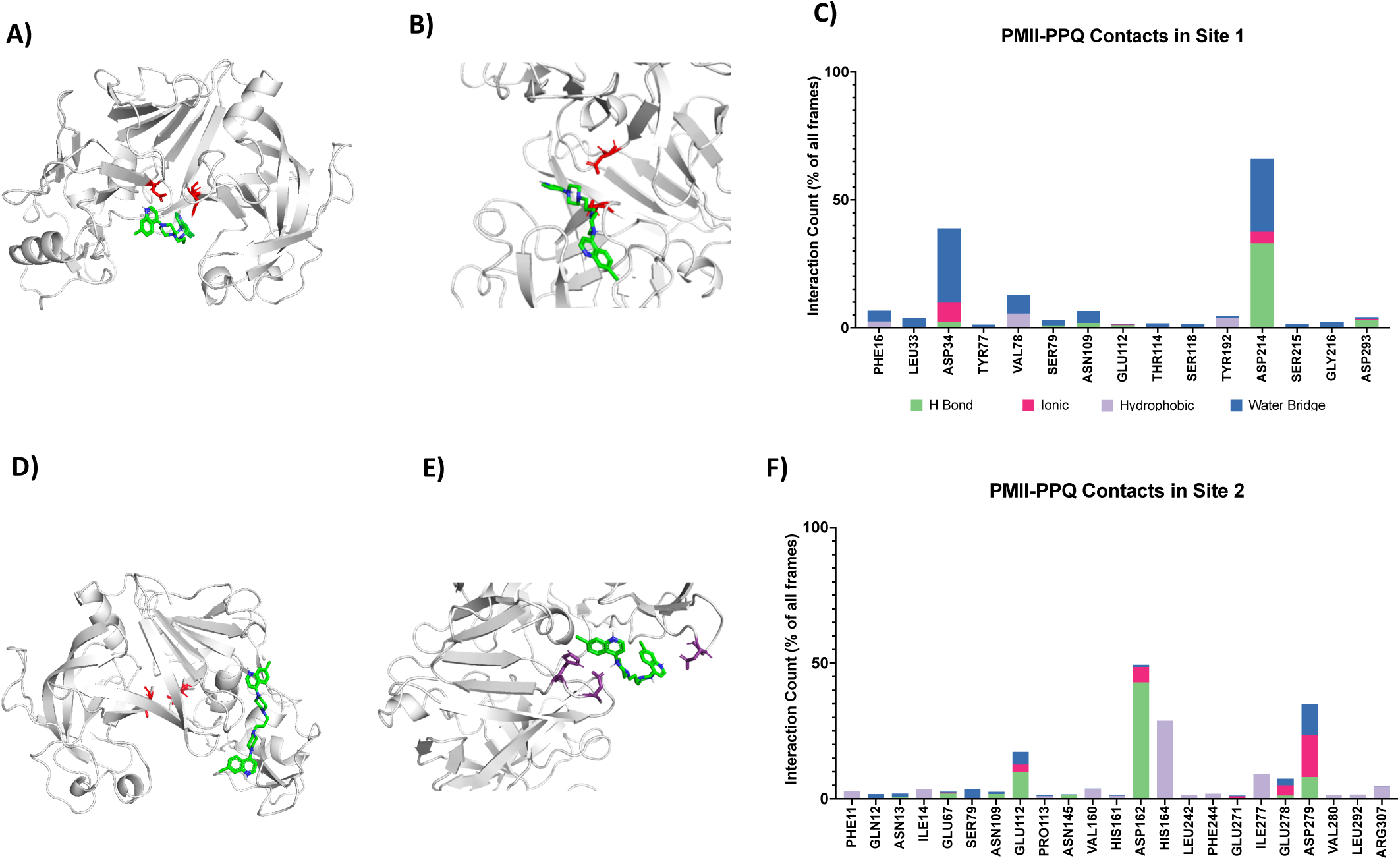
PMII energy minimized PPQ docked structures and side-chain interactions upon PPQ binding to two separate sites. A) Whole protein view of energy minimized XRMD_PMII (mature) with PPQ bound to site 1 (top left). B) Close-up view of energy minimized XRMD_PMII (mature) bound to PPQ at site 1 (top middle), catalytic Asp34 and Asp214 residues are shown in red. Nearly identical results are obtained with the AF2_PMII (mature) structure. C) Compiled side-chain interaction data for all residues associated with PPQ bound to site 1 for >1% of simulation time, Asp34 and Asp214 were identified as the most common (top right). D) Whole protein view of energy minimized XRMD_PMII (mature) with PPQ bound to site 2 (top right). E) Close-up view of XRMD_PMII with PPQ binding to site 2 (middle right) with Asp162, His164, and Asp279 residues shown in purple. F) Compiled side-chain interaction data for all residues associated with PPQ bound to site 2 for >1% of simulation time, Asp162, His164, and Asp279 were identified as the most common. Each side-chain interaction is shown as a stack representing 4 types of interaction: H Bond (green), ionic (pink), hydrophobic (purple), and water-bridged interaction (blue). The value for each residue interaction is the average of three independent 100 x ns energy minimization simulations.

**Table 1.** Predicted binding affinities from molecular docking simulations of the top three poses from each of three independent docking trials for PM II and PM III. . 8/9 poses for PM II localized to site 1 and one pose to site 2 whereas 9/9 of the highest affinity poses localized to site 1 in PM III. Lower binding energy indicates higher affinity binding.

| Trial #, Pose # | PM II | PM III |
| --- | --- | --- |
| Trial 1, Pose 1 | -8.8 kcal/mol (Site 1) | -8.1 kcal/mol |
| Trial 1, Pose 2 | -8.8 kcal/mol (Site 1) | -7.9 kcal/mol |
| Trial 1, Pose 3 | -8.5 kcal/mol (Site 1) | -7.9 kcal/mol |
| Trial 2, Pose 1 | -8.5 kcal/mol (Site 1) | -8.0 kcal/mol |
| Trial 2, Pose 2 | -8.1 kcal/mol (Site 2) | -8.0 kcal/mol |
| Trial 2, Pose 3 | -8.1 kcal/mol (Site 1) | -7.9 kcal/mol |
| Trial 3, Pose 1 | -8.6 kcal/mol (Site 1) | -7.9 kcal/mol |
| Trial 3, Pose 2 | -8.4 kcal/mol (Site 1) | -7.8 kcal/mol |
| Trial 3, Pose 3 | -8.2 kcal/mol (Site 1) | -7.9 kcal/mol |

For PM III (Fig. 6), although binding affinity is lower overall, all nine top poses from three independent docking simulations show PPQ localized to the catalytic cleft of the enzyme, positioned between His34 and Asp214 (Table 1; Fig. 5 C). Like PM II, we see a lower affinity secondary site (2/27 total poses from three independent docking simulations). Together, these results indicate that PPQ likely occupies the catalytic clefts of PM II and PM III near catalytic residues, supporting the possibility of enzyme inhibition under high-dose conditions. Predicted binding affinities for the highest-ranked docking poses ranged from -8.1 to -8.8 kcal/mol for PM II and from -7.9 to -8.1 kcal/mol for PM III (Table 1), consistent with energetically favorable interactions between PPQ and both plasmepsins and a slightly higher predicted binding affinity for PM II than PM III.

**Figure 6.**
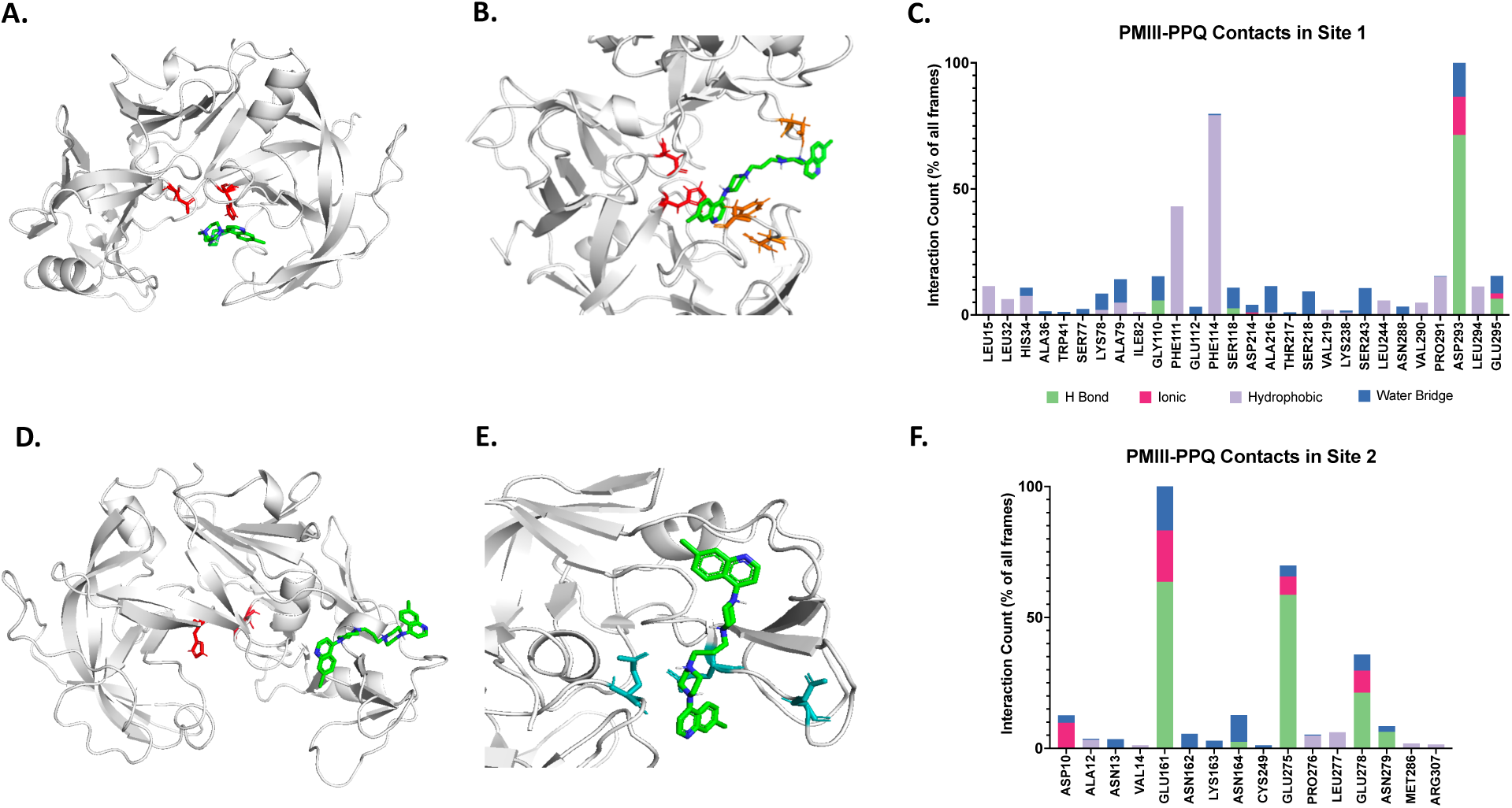
PM III energy minimized PPQ docked structures and side-chain interactions upon PPQ binding to two separate sites. A) Close-up view of energy minimized AFMD_PMIII (mature) bound to PPQ at site 1 (top left), catalytic His34 and Asp214 residues are shown in red and Phe11, Phe114, and Asp293 residues are shown in orange. B) Close-up view of AFMD_PMIII with PPQ binding to site 2 (top right) with Glu161, Glu275, and Glu278 residues shown in teal. C) Compiled side-chain interaction data for all residues associated with PPQ bound to site 1 for >1% of simulation time, Phe111, Phe114, and Asp293 were identified as the most common interactions. D) Compiled side-chain interaction data for all residues associated with PPQ bound to site 2 for >1% of simulation time Glu161, Glu275, and Glu278 were identified as the most common. Each side-chain interaction is shown as a stack representing 4 types of interaction: H Bond (green), ionic (pink), hydrophobic (purple), and water-bridged interaction (blue). The value for each residue interaction is the average of three independent 100 x ns energy minimization simulations.

### Molecular Dynamics Simulation of PPQ bound to PM II

To further characterize the predicted PPQ binding sites, we assessed whether PPQ forms stable interactions with residues involved in substrate recognition and catalysis using molecular dynamics simulations. After docking each protein structure to PPQ in three separate trials, the highest affinity pose for each site underwent energy minimization (Figs. 5 A, B; 6 A, B). The resulting trajectories were merged and clustered to create averaged docked structures that reveal molecular contacts, and simulation interaction diagrams between protein and ligand were generated in Maestro (Fig 5 C,D and Fig 6 C,D). In site 1 for PM II, interactions between PPQ and Asp34 and Asp214 were present for approximately 39% and 66% of simulation time, respectively (Fig. 5 A, C). This indicates that PPQ is likely associating with PM II in the active site via interaction with catalytic residues that are the most important for Hb proteolysis. In the secondary site for PM II, interactions between PPQ and Asp162, His164, and Asp279 were present for approximately 50%, 29%, and 35% of simulation time, respectively. In the catalytic site for PM III (comparable to site 1 for PM II), interactions between PPQ and Phe111, Phe114, and Asp293 were present for approximately 43%, 80%, and 100% of simulation time, respectively, whereas interaction with His34 that replaces Asp34 found in PM II was ∼10 % of simulation time and interaction with Asp214 was < 5 % of simulation time compared to 66 % for this residue in PM II. In the secondary site for PM III (comparable to site 2 for PM II), interactions between PPQ and Glu161, Glu275, and Glu278 were present for approximately 100%, 70%, and 36% of simulation time, respectively. Collectively, these data suggest that PPQ binds within the catalytic domains of both PM II and PM III domains but that molecular contacts for docking differ for the two related enzymes resulting in altered binding affinity.

## Discussion

The bimodal PPQ dose-response phenotype represents a distinct and poorly understood form of antimalarial drug resistance that cannot be explained by conventional measures of drug susceptibility. Although amplification of *pm II/III* has been consistently associated with this phenotype, the biological mechanism linking increased *pm II/III* copy number to enhanced survival at high PPQ concentrations remains unresolved. In this study, we sought to develop a more comprehensive understanding of the molecular changes linking *pm II/III* copy number variation to the bimodal PPQ survival phenotype Our findings support a new mechanistic framework in which increased PM II/III abundance, resulting from *pm II/III* amplification, enables direct PPQ-plasmepsin interactions while preserving sufficient proteolytic capacity to maintain Hb metabolism under high PPQ exposure. We reached this proposed model by integrating phenotypic, transcriptional, metabolomic, and *in silico* structural analyses across isogenic PPQ-resistant parasites differing only in *pm II/III* copy number..

We investigated the influence of parasite stage and exposure duration on PPQ dose-response curve dynamics. Consistent with previous findings, ring- and trophozoite-stage parasites carrying multiple copies of *pm II/III* display increased survival under high PPQ concentrations (10,16,17). However, parasites initially exposed to PPQ as schizonts, regardless of underlying *pm II/III* copy number, no longer exhibited this secondary survival response. Stage-dependent variation in parasite survival following PPQ exposure has previously been described (16), although that study did not specifically examine late schizont-stage parasites or evaluate the effects in the context of bimodal dose-response curve shape. Interestingly, this stage-specific effect altered parasite survival only at high PPQ concentrations and did not substantially shift the overall resistant phenotype as measured through limited-point IC_50_ assays or PSA. These findings indicate that the contribution of *pm II/III* amplification to the bimodal survival phenotype is distinct from the broader PPQ resistance conferred by *pfcrt*. Instead, *pm II/III* copy number likely modulates parasite survival under high-dose PPQ exposure rather than altering baseline drug susceptibility.

The persistence of this phenotype following shortened PPQ exposures further suggests a critical role of the developmental stage at which parasites encounter drug, rather than exposure duration. One possible hypothesis for this stage specificity is the reduced transcriptional expression of *pm II/III* during late schizogony, which may negate the effect of carrying multiple genomic copies of these genes. Importantly, PPQ exposure itself did not induce changes in expression of *pm II/III.* While external stressors such as heat shock have been shown previously to influence transcriptional regulation of these genes (27), our data indicate that the bimodal survival phenotype is not driven by acute transcriptional changes of *pm II/III* in response to increasing PPQ concentrations, but by differences in PM II/III abundance within the digestive vacuole resulting from genomic copy number differences.

The contribution of *pm II/III* amplification to the bimodal PPQ survival phenotype is highly dependent on the broader PPQ-resistant background established by *pfcrt* genotype. Numerous studies have identified PfCRT as the primary determinant of PPQ resistance, with novel PfCRT mutations (T93S, H97Y, F145I, I218F, M343L, G353V, and G367C) promoting the active transport of PPQ out of the digestive vacuole and thereby reducing intravacuolar drug accumulation (2,6,13,28–32). However, as has been observed for PfCRT-mediated resistance to multiple aminoquinoline antimalarials, altered transport activity is also associated with broader changes to digestive vacuole physiology and homeostasis (13,33,34).

Under this framework, the epistatic relationship between *pfcrt* and *pm II/III* may reflect either a requirement for reduced PPQ accumulation within the digestive vacuole or broader alterations to the digestive vacuole microenvironment that permit increased plasmepsin abundance to influence parasite survival under high PPQ exposure. Overexpression of *pm II/III* in the PPQ-sensitive Dd2 background does not independently lead to a bimodal survival response to PPQ, whereas this phenotype is recovered upon the disruption of PfCRT through the addition of verapamil (35). Similarly, experimental perturbation of digestive vacuole proton homeostasis through concanamycin A or CCCP treatment significantly reduced the magnitude of the bimodal PPQ survival response, despite PPQ itself having minimal direct effects on digestive vacuole pH (17). Together, these findings suggest that the contribution of *pm II/III* amplification is closely tied to broader PfCRT-driven changes in digestive vacuole homeostasis rather than functioning as an isolated determinant of PPQ survival. Interestingly, copy number variation has now been associated with multiple members of the plasmepsin family. In addition to *pm II/III* amplification in PPQ resistance, recent studies of resistance to *plasmepsin IX* and *plasmepsin X* inhibitors identified both point mutations and copy number amplification involving both of these genes, suggesting that modulation of plasmepsin abundance may represent a broader adaptive mechanism within the plasmepsin family (36).

Given the central role of PM II/III in Hb catabolism within the digestive vacuole (23,37), and previous reports linking aminoquinoline resistance-conferring PfCRT variants to reduced transport of Hb-derived peptides (33,38–40), we examined metabolic responses to PPQ exposure across isogenic parasites differing only in *pm II/III* copy number within a shared PPQ-resistant *pfcrt* background. Under 2 µM PPQ, which was used in the metabolomics experiments as a scaled equivalent of concentrations associated with the secondary survival phase, parasites with multiple copies of *pm II/III* exhibited increased abundance of multiple peptides previously associated with Hb degradation relative to the single-copy parasite FG0305. Both the number of altered peptides and the magnitude of peptide accumulation increased with *pm II/III* copy number. These changes were specific to high PPQ exposure and were not observed under untreated conditions or lower PPQ concentrations, indicating that these metabolic perturbations emerge under conditions associated with the bimodal survival phenotype.

Notably, previous studies reported increased abundance of several Hb-derived metabolites, including PVNF, PE, PD, DLH, and VD, in parasites carrying resistance-associated *pfcrt* mutations that actively transport aminoquinoline antimalarials from the digestive vacuole (13,33,34,38,41). Because all parasites examined in this experiment share the same PPQ-resistant pfcrt background, the progressive increase in both the abundance and diversity of these metabolites with increasing *pm II/III* copy number is not due to differences in PfCRT genotype alone. Instead, we propose that active transport of PPQ by mutant PfCRT contributes to the accumulation of intermediate products of Hb degradation within the digestive vacuole. Meanwhile, increased Hb digestion resulting from increased PM II/III abundance within the digestive vacuole may increase the production of these same intermediates, leading to the progressive increase in both the abundance and diversity of metabolites observed with increasing *pm II/III* copy number. Together, these findings provide a potential metabolic explanation for the observed epistatic relationship between *pfcrt* and *pm II/III*, whereby independent effects of mutant PfCRT and increased plasmepsin abundance converge on enhanced digestive vacuole metabolism under the same high PPQ conditions associated with increased parasite survival.

While the metabolomic findings point to an increased PM II/III abundance contributing to enhanced digestive vacuole metabolism under high PPQ exposure, molecular modeling also supports a mechanism of direct interactions between PPQ and PM II/III proteins. PPQ consistently mapped to two recurrent regions of PM II, including the catalytic cleft containing the active-site aspartates and a secondary pocket formed by the L3–L4 loop. Importantly, both regions contain residues previously implicated in substrate binding and processing, with the L3–L4 pocket specifically proposed to participate in substrate anchoring during Hb cleavage (26,42). Similar localization patterns were also observed for PM III. While these analyses do not establish direct biochemical binding, they suggest that PPQ may interact with previously characterized substrate-binding regions of plasmepsin proteins. Additional studies using purified plasmepsin proteins and direct biochemical binding assays will be required to validate these predicted interactions and determine whether they contribute to altered PPQ susceptibility.

Under this framework, *pm II/III* amplification increases the total plasmepsin pool within the digestive vacuole. This expanded plasmepsin pool may allow a subset of PM II/III molecules to interact directly with PPQ while sufficient active enzyme remains available to sustain Hb catabolism, a balance that may not be achievable in parasites carrying fewer copies of *pm II/III.* This framework also provides a potential explanation for the stage specificity of the bimodal phenotype, as the marked reduction in *pm II/III* transcription during schizogony would decrease the total plasmepsin pool available within the digestive vacuole, thereby reducing the opportunity for PPQ-plasmepsin interactions despite persistent differences in genomic copy number. This model may also explain the concentration dependence of the bimodal phenotype, whereby increasing PPQ concentrations progressively favor PPQ-plasmepsin interactions, allowing the effects of *pm II/III* amplification to emerge only under high-dose conditions. Importantly, previous work showed that deletion of *pm II* and *pm III* in combination produces minimal fitness defects under nutrient-sufficient conditions, suggesting that partial disruption of plasmepsin function may be biologically tolerated during asexual blood-stage growth (43). Consistent with this interpretation, deletion of *pm II* and *pm III*, both individually and in combination, in a multicopy *pm II/III* parasite background was shown to slow Hb digestion and parasite development while remaining compatible with parasite survival (17). Increased plasmepsin abundance may therefore permit a subset of PM II/III proteins to interact with PPQ while maintaining sufficient proteolytic activity to support continued Hb catabolism within the digestive vacuole. Together, these findings provide a potential mechanistic framework linking *pm II/III* amplification to the bimodal PPQ survival phenotype.

Despite the current overall success of DHA+PPQ in Africa, multiple studies have reported duplications of *pm II* in clinical isolates from both East and West Africa (44–46). In some cases, duplications were found in >30% of samples in Burkina Faso and Uganda (46). While none of the major PPQ resistance conferring PfCRT substitutions have yet to been found in this region, the co-occurrence of African samples with *kelch13* mutations and duplications of *pm II* threatens the future efficacy of DHA+PPQ in Africa (46). Here, we applied a comprehensive experimental framework to investigate the relationship between *pm II/III* copy number amplification and the bimodal PPQ survival phenotype through targeted manipulation of PPQ exposure conditions, transcriptional and metabolomic analyses, and structural modeling of potential PPQ–plasmepsin interactions. Although further validation will be required to fully define the molecular mechanisms underlying this phenotype, our findings demonstrate that the bimodal survival response is influenced not only by *pm II/III* genomic copy number, but also by stage-specific differences in plasmepsin expression across the parasite life cycle. Together, these findings provide important insight into how increased plasmepsin abundance contributes to parasite survival under high PPQ exposure and further refine our understanding of the mechanisms linking *pm II/III* copy number amplification to the bimodal PPQ resistance phenotype in *P. falciparum*.

## Materials and Methods

### Parasites used in this study

Seven *P. falciparum* parasites representing distinct combinations of *pfcrt* genotype and *pm II/III* copy number were used to investigate the relationship between *pm II/III* amplification and PPQ response. KH004, a Cambodian PPQ-resistant isolate carrying a PPQ-resistant *pfcrt* allele (Dd2 + G367C) and four copies of *pm II/III*, served as the primary parental background for this study. FG0305 is a selfed version of KH004 recovered from the KH004×Mal31 genetic cross and is isogenic to KH004 except at the *pm II/III* locus, where it has only a single copy of these genes. Additionally, two isogenic clones of KH004, KH004-D9 and KH004-E4, were utilized. These clones were recovered through limited dilution cloning of KH004 following a 48-h selection with 125 nM PPQ. These closes are isogenic to the KH004 parental parasite but have 5 and 3 copies of *pm II/III* respectively. Finally, Mal31 was used as a PPQ-sensitive control. This parasite was a parent of the KH004×Mal31 genetic cross and has a PPQ-sensitive 3D7-like *pfcrt* genotype as well as a single copy of *pm II/III*.

### *Plasmepsin II/III* copy number determination

*Plasmepsin II/III* copy number was estimated using sequencing read depth across an extended 67-kb region on chromosome 14 (positions 260,000–327,000), encompassing the tandemly duplicated *pm II* and *pm III* genes as well as 28 kb of flanking sequence upstream of *pm I* and downstream of *pm III*. Read coverage at each genomic position was calculated from deduplicated BAM files using *bedtools* (https://bedtools.readthedocs.io/en/latest/). Coverage profiles were visualized using Integrative Genomics Viewer (IGV) to confirm duplication boundaries and consistency across samples. Copy number was calculated as the ratio of mean read coverage across the duplicated *pm II/III* repeat unit to mean coverage across the non-duplicated flanking regions. This normalization accounts for variation in overall sequencing depth and enables direct comparison of copy number across parental and progeny parasites.

### Parasite Culture

Cryopreserved parasites were thawed and cultured in complete media consisting of 0.5% Albumax II (Gibco, Life Technologies) supplemented RPMI 1640 with L-glutamine (Gibco, Life Technologies) with additional 50 mg/L hypoxanthine (Calbiochem, Sigma-Aldrich), 25 mM HEPES (Corning, VWR), 10 µg/mL gentamycin (Gibco, Life Technologies), and 0.225% sodium bicarbonate (Corning, VWR). Parasite cultures were maintained at 5% hematocrit in O+ red blood cells (Interstate Blood Bank, Memphis TN) in separate flasks and maintained at consistent pH (7.0-7.5), temperature (37°C), and atmosphere (5% CO_2_/5% O_2_/90% N_2_). Culture media was changed every 48 hours, corresponding to one intraerythrocytic development cycle. Parasitemia and parasite stage were assessed daily through both microscopy and flow cytometry, with parasitemia being kept below 2%.

### Drug susceptibility assays

All assays were performed with synchronized parasites following a single layer Percoll separation. 500µL of infected erythrocytes, > 1% parasitemia and >50% schizonts, were packed and suspended in 2mL of RPMI and layered over a 70% Percoll (Sigma-Aldrich) layer in 1x PRMI and 13.3% Sorbitol in phosphate buffered saline (PBS). Layered cultures were centrifuged for 10 min at 1575×g with no brake. The top layer of purified schizonts were removed, washed, and resuspended in complete media and uninfected red blood cells at 5% hematocrit. Parasites were set up in assays as either early rings (6 hours post synchronization (hps)), late rings (20 hps), trophozoites (32 hps), or schizonts (44 hps) depending on assay specifications. One hour prior to assay setup, parasitemia and stage were determined via flow cytometry. Parasites were stained with a combination of SYBR Green I and Syto61 and 50,000 events were counted on a Guava easyCyte HT (Luminex Co.) flow cytometer. Drug-susceptibility assays were performed on cultures in which over 80% of parasites were at the intended stage.

For generating dose-response curves, parasites were diluted to 0.15% parasitemia and 2% hematocrit and set up in a 96-well plate with two technical replicates at a volume of 150µL per well. To capture a broad range of responses, parasites were tested against 20 unique PPQ concentrations derived from two 10-point serial dilutions, starting at 3 µM and 500 nM, alongside untreated and uninfected red blood cell controls. Parasites received consistent PPQ exposure for 72 hours unless otherwise noted. In instances where duration of exposure was limited, parasites were washed 3x with RPMI and returned to untreated complete media for the remained of the 72 hours. Parasite density was determined through SYBR Green fluorescence and dose response curves were generated by plotting percent survival against the log of drug concentration.

PSA was conducted as previously described in Kane et al. 2024 (10). Briefly, synchronized early ring stage parasites were exposed to a single 200 nM dose of PPQ for 48 hours. Parasites were then washed with RPMI and returned to drug-free complete media before collecting samples for analysis at the 120 h timepoint. Parasite prevalence was determined via qPCR, and relative survival was calculated as fold change between treated and untreated samples for three technical replicates (three replicates within the plate as well as three independent qPCR reads).

### Assessment of *pm II/III* transcriptional expression

All transcriptional analysis was performed using synchronized parasites at one of the intraerythrocytic developmental timepoints listed above. In the case of transcriptional assessment following PPQ exposure, protocols were based on previously optimized work of Siwo et al. 2015 (47). Total RNA isolation was performed using a TRIzol-chloroform extraction. 5 volumes (v/v) of Trizol (Invitrogen) were added to infected packed red blood cells and thoroughly homogenized. Chloroform was added at 1:5 ratio to the TRIzol mixture, vortexed, and centrifuged at 9,000 × g for 20 minutes to separate phases. The supernatant containing total RNA was removed and mixed with an equal volume of 100% molecular grade ethanol (ThermoFisher). RNA was extracted and purified using RNA Clean & Concentrator-5 kit (Zymo) following manufacturer’s instructions. RNA was eluted in 25 µL of DNAse/RNAse free water, and purity and concentration were assessed through Nanodrop. Purified RNA was treated with DNAse I (Invitrogen) for 10 minutes prior to cDNA synthesis via Superscript III (Invitrogen). cDNA was treated with 2 units RNAse H (Invitrogen) for 20 minutes to remove DNA-RNA hybrids. Synthesized cDNA concentration was assessed through Nanodrop (ThermoFisher).

Analysis of *pm II/III* expression was performed by RT-qPCR. 100 ng of pure cDNA was used per well. Each cDNA sample was run in triplicate using primers specific to *pm I pm II*, *pm III*, and *actin1* for 40 cycles using the following protocol: denaturation at 95°C for 20 seconds, followed by 40 cycles of 95°C for 1 second, 62.5°C for 30 seconds, and 65°C for 15 seconds. The qPCR was set to “fast” mode, reaction volume is 10 μl and set to “SYBR detector no quencher”. All samples were normalized to *actin1* (ΔCt), and expression was presented as a fold change (2^ΔCt^) relative to *actin1*. In the case of PPQ treated samples, all samples were initially normalized to *actin1*, before as second normalization to the untreated condition of that same parasite (ΔΔCt) and presented as fold change relative to the untreated expression level (2^ΔΔCt^) within that individual parasite.

### Metabolite profiling

Treatment of parasites and extraction of metabolites were performed based on methods described by Allman et al. 2016 (24). Briefly, 400 mL of synchronized trophozoite-stage culture (4% hematocrit and ∼5% parasitemia) was magnetically purified using a SuperMACS II magnet and MACS CS columns. Following a 1-hour recovery period, parasites were exposed to one of five treatment conditions: untreated control, 200 nM PPQ, 500 nM PPQ, 2 µM PPQ, or 10 nM atovaquone (ATQ) for 2.5 hours. These conditions were selected based on prior work demonstrating that short exposures at concentrations approximately ten times the parasite IC_50_ produce strong metabolic signals.

The selected PPQ concentrations represent 10× the concentration corresponding to the lowest parasite survival prior to the secondary phase of the dose-response curve (500 nM), the concentration at the peak of the secondary survival phase (2 µM), and the standard concentration used in the piperaquine survival assay (200 nM). To facilitate direct comparison between parasite lines and treatment conditions, all isogenic PPQ-resistant parasites were processed together within the same experimental batch. Following the 2.5 h exposure, parasites were washed with cold phosphate-buffered saline (PBS) and metabolism was quenched with ice-cold 90% methanol containing 0.5 µM ^13^C_4_,^15^N-aspartate as an internal standard.

All samples were processed in biological triplicate with method blanks included to account for background signal. Samples were randomized prior to analysis, and pooled quality control samples and blanks were run regularly throughout the analytical sequence to monitor instrument performance and reduce technical variation. Prior to analysis, samples were dried under N_2_ gas and resuspended in either 50 µL or 80 µL of 0.5 µM ^13^C_5_,^15^N-glutamate in 90% methanol depending on total parasite concentration for a final 10^6^ former cells/μL concentration. A 10 µL aliquot of each sample was injected into a Thermo Scientific Vanquish Flex UHPLC for separation of metabolites using a Waters XBridge BEH Amide Column (catalog number: 186004861). Chromatographic separation was carried out using a 25-minute HILIC method gradient of 5% acetonitrile (A) and 90% acetonitrile (B) both supplemented with 20mM of ammonium hydroxide and ammonium acetate. The resulting eluate was sent into a Thermo Scientific Orbitrap Exploris 120 mass spectrometer for analysis. Following data acquisition, the raw files were converted into .mzML files using MSConvert (48). Peak detection, alignment, and metabolite annotation were performed using El-Maven (49) with the Llinás laboratory curated metabolite reference library. The peak areas were corrected using the ^13^C_4_, ^15^N-aspartate signal and blank signals were subtracted. Metabolites were filtered using pooled quality control samples, and those with a relative standard deviation greater than 30% were excluded. In total, 123 targeted metabolites passed quality control and were retained for downstream analysis.

### Protein Structure Preparation

We first imported the atomic-resolution structure of PM II from *P. falciparum* solved by x-ray diffraction (PDB code: 1LF3 (50)) into Maestro (51). The structure was prepared using Protein Preparation Wizard where bond orders were assigned and protonation states of ionizable residues were fixed at pH 5.0, near the pH of the DV where PMII is found, with ProtAssign (52). For PM III, the same was done except the structure was obtained from the AlphaFold2 database (AFDB: Q8IM15F1).

### Molecular Dynamics

Restrained energy minimization with heavy atoms converged to 0.30 Å was performed using the OPLS4 force field. Replica Exchange MD (REMD) energy minimization was performed either with or without docked drug. In both cases, the Maestro System Builder Panel was used to neutralize charges with sodium and chloride ions and solvate the protein with simple point-charge (SPC) water within an orthorhombic box expanding 10 Å beyond the protein in the X, Y, and Z dimensions. REMD was performed using Desmond run within Maestro (53,54). Each calculation began with a default system relaxation protocol followed by simulation in an isothermal, isobaric NPT ensemble with constant particle number (N), pressure (P; 1.01325 bar), and temperature (T; 310K). Simulation interaction diagrams were then used to analyze resultant protein−drug interactions. Each simulation was run for 100 ns. Prior to docking, the time step used is 10 ps and the latter half of three independent simulations each with randomized starting velocity, were used to generate an energy-minimized structure. This structure was used for subsequent docking studies and further REMD with drug present; each docked structure underwent three independent simulations each 100 ns with 100 ps time steps. All frames were used for analysis of the minimized docked structures.

### Drug Docking

AutoDock Vina was used to dock PPQ to PMII and PMIII (55). In brief, REMD minimized structure was prepared using AutoDock MGL Tools where waters were deleted, polar hydrogens and Kollman Charges were added, and the protein was saved as a PDBQT. To prepare the drug molecule the SDF file was obtained from PubChem. The file was imported into PyMOL v2.4.0 (Schrödinger, LLC, New York, NY) to manage protonation states and then exported as a PDB file. The drug file was then imported into AutoDock MGL Tools, prepared, and saved as a PDBQT file. Grid boxes encompassing the whole protein were used. The binding simulation was run in the command window, and the outputs were analyzed using PyMOL. Three independent docking simulations were run, and the top poses in each site were used in further analysis.

## Data Availability

Sequences for all parasites used in this study can be found on the NCBI Sequence Read Archive (SRA, https://www.ncbi.nlm.nih.gov/sra) under the project number PRJNA524855. All code used for sequence, metabolite, and transcript analysis are available through GitHub (https://github.com/FerdigLab/PPQ-Bimodal-Response). Additional data related to this work may be requested from the authors.

## Acknowledgements

This work was supported by the National Institutes of Health (NIH) program project grant P01 AI127338 (to M.T.F.). The parental line, Mal31-9040-C11, used in the Mal31 × KH004 cross was sampled from a Malawian patient in 2016 as part of a cross-sectional study funded by the Wellcome Trust of Great Britain (Grant no. 099992/Z/12/Z) and provided by Standwell Nkhoma.

We thank the TRAC II Collaboration, Rob van der Pluijm and Arjen Dondorp, for providing KH004-020-019-H9. We thank the patients who provided parasites used in this work.

