## Supplemental Figures for "A Multidimensional Analysis of the Bimodal Piperaquine Response in *Plasmodium falciparum*"

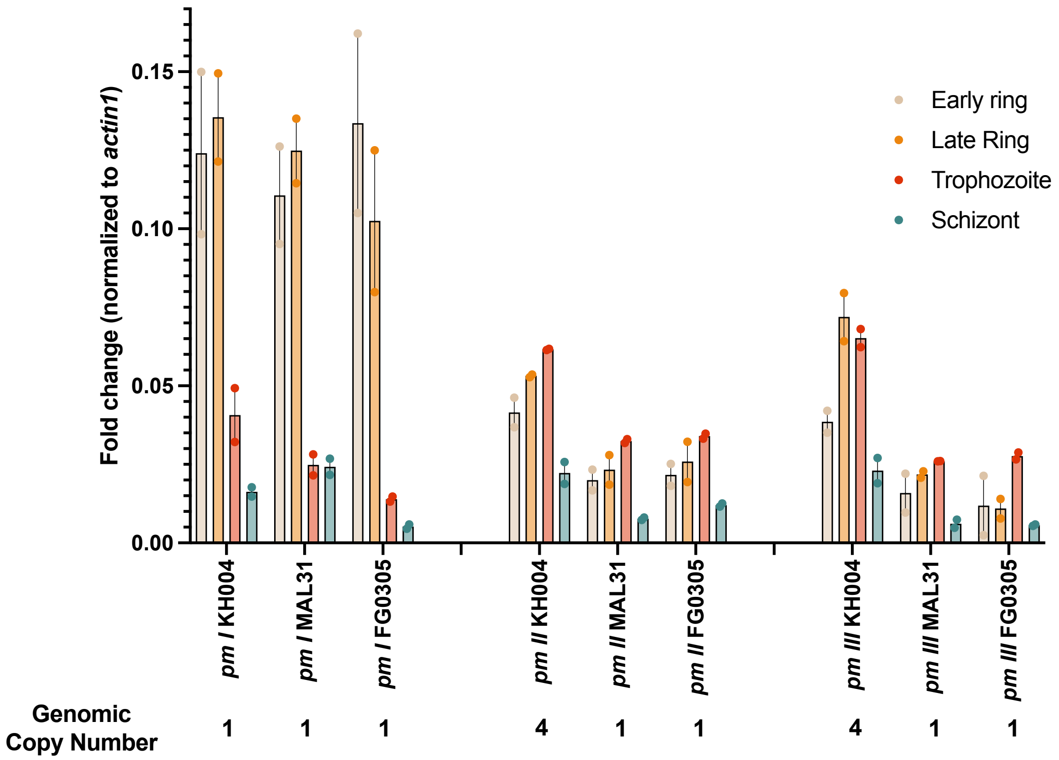


**Supplementary Fig. 1.** Expression of *plasmepsin* genes across the asexual lifecycle. Expression of the DV localized *plasmepsin* genes is variable across the asexual parasite lifecycle. *Pm I* is expressed at the highest levels early in the parasite lifecycle with steady decrease in expression as the parasite develops. *Pm II* and *pm III* are most highly expressed during trophozoite stage, when the parasite is metabolizing the bulk of hemoglobin.

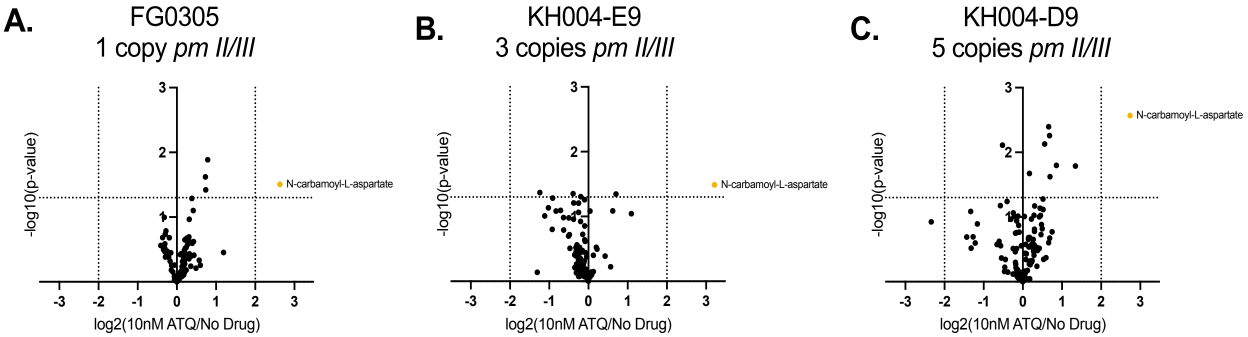


**Supplementary Fig. 2. Atovaquone control confirms consistent metabolically active response across parasite lines.** Volcano plots comparing atovaquone (ATQ)-treated samples to no-drug controls across all parasite lines show a consistent increase in N-carbamoyl-L-aspartate across all parasite backgrounds. The increase in this metabolite is consistent with a metabolic response to ATQ, which disrupts pyrimidine biosynthesis.

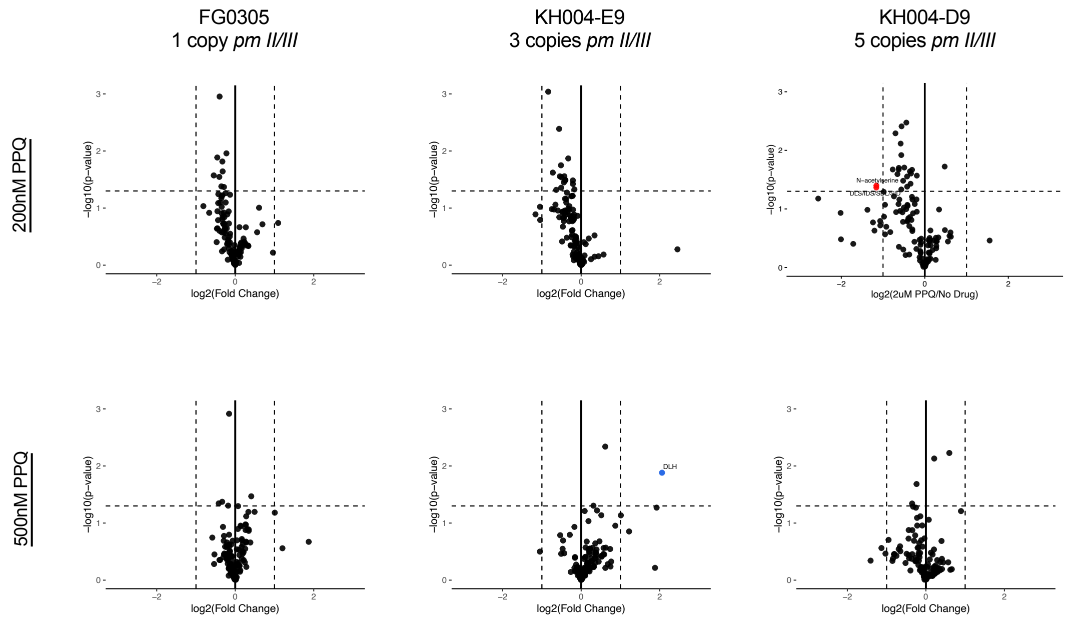


**Supplementary Fig 3. Metabolic response following lower-dose PPQ exposure.** Volcano plots showing differences in metabolite abundance relative to untreated controls following exposure to 200 nM (top row), 500 nM (bottom row) PPQ. Columns from left to right correspond to KH004-D9 clone (5 copies of *pm II/III*), KH004-E9 clone (3 copies of *pm II/III*), and FG0305 (1 copy of *pm II/III*). Each point represents a detected metabolite plotted as $\log_2$ fold change relative to untreated parasites (x-axis) versus $-\log_{10}$ (p-value) (y-axis). Vertical dashed lines indicate fold-change thresholds and the horizontal dashed line indicates the significance threshold. Data points represent the mean abundance across three technical replicates within a single biological replicate. Across all parasites and conditions, there was a minimal significant difference in metabolite abundance compared to untreated samples. The only observed metabolite to be significantly increased in the 3 copy KH004-E4 clone following 500 nM PPQ selection. Additionally, N-acetylserine and a set of tripeptides were found at decreased abundance in KH004-D9 following 200 nM PPQ.

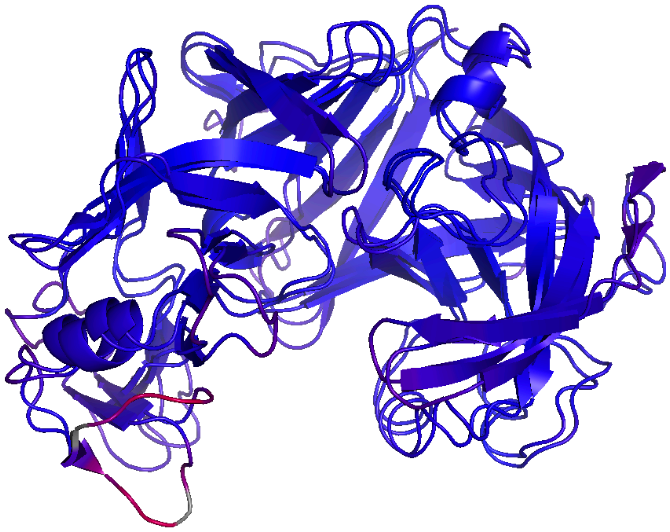

**Supplementary Fig 4: All atom RMSD superposition of minimized mature XRMD_PMII vs minimized mature AFMD_PMII structures.** Results are color coded Minimum Distance (blue): 0.12 Maximum Distance (red): 19.75, average distance = 2.36 Å, showing x-ray and AF2 protein structures for these enzymes are highly similar.

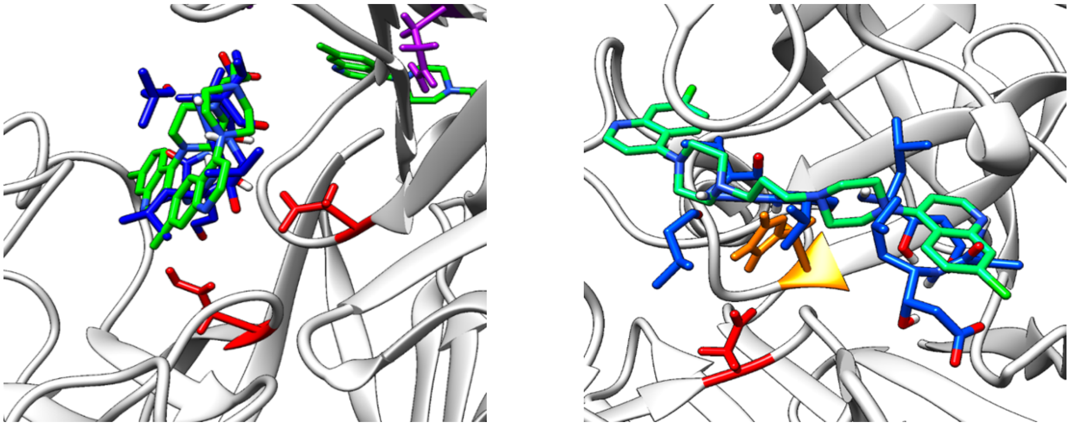

**Supplementary Fig. 5. Predicted PPQ binding sites overlap with known plasmepsin inhibitor binding regions.** Molecular docking simulations identified the catalytic cleft of both PM II (left) and PM III (right) as a recurrent interaction site for PPQ (green). Additional docking simulations using the aspartic protease inhibitor Pepstatin A (blue) demonstrated substantial overlap between the predicted binding poses of the two ligands within the active-site region of each enzyme. In both PM II and PM III, PPQ and Pepstatin A localized adjacent to the catalytic Asp (red) and His (orange) residues within the protein active sites, consistent with interactions occurring within previously characterized substrate-binding regions of the protease.
